# Intra-host quasispecies reconstructions resemble inter-host variability of transmitted chronic hepatitis B virus strains

**DOI:** 10.1101/2023.05.15.540814

**Authors:** L.S. Le Clercq, S.M. Bowyer, S.H. Mayaphi

## Abstract

The hepatitis B virus is a partially double stranded DNA virus in the Hepadnaviridae family of viruses that infect the liver cells of vertebrates including humans. The virus replicates through the reverse transcription of an RNA intermediate by a viral polymerase, akin to retroviruses. The viral polymerase has high replication capacity but low fidelity and no proofreading activity resulting in a high mutation rate. This contributes to the emergence of a cloud of mutants or quasispecies within host systems during infection. Several host and viral factors have been identified that contribute to mutations and mutation frequency in shaping viral evolution, however, because the dynamics of viral evolution cannot be understood from the fittest strain alone, the need exists to sequence and reconstruct intra-host diversity, recently made possible through next generation sequencing. Due to the extensive pipeline of bioinformatic analyses associated with next generation sequencing studies are needed to ascertain if quasispecies reconstruction methods and diversity measures accurately model known diversity. Here, next generation sequencing and various quasispecies reconstruction methods are used to model the natural evolution of viral populations across the full genome of hepatitis B virus strains from South Africa. This study illustrates that (i) different methods of quasispecies reconstruction reconstruct the same amount of diversity, (ii) intra-host diversity derived from full quasispecies analyses resembles diversity measures obtained from previous methods, (iii) inter-host diversity resembles the diversity between closely related quasispecies variants, (iv) diversity is increased in HIV-negative individuals, and (v) corroborate that seroconversion of HBV biomarkers increases mutation rates.

## Introduction

The Hepatitis B Virus (HBV), Orthohepadnavirus homonensis, is a deoxyribonucleic acid (DNA) virus that infects humans, first discovered in 1965 by Blumberg [5, 6]. It is one of at least 18 species within the Hepadnaviridae family which has five genera, two of which are well characterised (Ortho- and Avihepadnaviridae) and three (Meta-, Herpeto-, and Parahepadnaviridae) which have only been recognised recently: with new members still being discovered [44, 32, 50]. These viruses infect the hepatocytes of the liver in the five most common classes of vertebrates: mammals, birds, fish, reptiles, and amphibians. HBV is specifically classified in the Orthohepadnavirus genus of DNA viruses infecting the liver of mammals, including the related viruses that infect many non-human primates as well as those infecting squirrels, woodchucks, bats, and equines. Studies have sub classified HBV into nine widely accepted genotypes or viral subspecies, denoted A to J, based on pair-wise differences of more than eight and less than seventeen percent across the full genome. These genotypes are further partitioned—with the exception of genotypes E, G and H—into subgenotypes or molecular subtypes, based on pairwise differences larger than four percent [42, 53]. HBV genotypes seem to follow a remarkedly conserved geographic distribution, reviewed by Kramvis et al. [28].

HBV is primarily a blood-borne virus that is transmitted between individuals through direct contact of bodily fluids such as blood but can also be transmitted sexually [23] through other body fluids. Exposure can either result in an acute infection, where the host system eventually clears the infection and recovers, or may result in a chronic and persistent infection [55]. Upon exposure, the virus infiltrates host liver cells and initiates replication, protein transcription, and new virion formation. This occurs through the epigenetic regulation [8], transcription and translation of viral open reading frames from covalently closed circular HBV genome copies (“mini-chromosomes”) in the nucleus; including the core gene, surface gene, X gene, and viral polymerase. The HBV genome is replicated by the reverse transcription of a viral ribonucleic acid (RNA) intermediate, the pre-genomic RNA, following encapsidation in the cytoplasm [35, 26]. Several viral components are detectable in the blood of infected individuals and have been developed into biomarkers of infection and disease progression; reviewed by Kramvis et al. [29]. Copies of HBV DNA, linearised during the reverse transcription of pre-genomic RNA, can also be incorporated into the host genome [53]. The integration of hepatitis viruses into host genomes may not maintain a chronic infection, however, it was recently illustrated in several passerine bird species, including finches, that parts of the viral DNA may become endogenous to the organism it infects and remain in their genomes long after their species are no longer common hosts [17].

RNA viruses such as the hepatitis C virus and influenza virus, and reverse transcriptase dependent viruses such as HBV and the human immunodeficiency virus (HIV), show a high degree of intra-host variability. This is likely due to the high replication capacity yet low fidelity and lack of proofreading activity of viral polymerases; typically intro-ducing between 1.0 *×* 10^*−*5^ and 1.0 *×* 10^*−*3^ substitutions per site per cycle in retroviruses [10]. The substitution rate, calculated per site per year, for HBV has been approximated to be between 1.4 *×* 10^*−*5^ and 5 *×* 10^*−*5^: comparable to the rate in retroviruses but nearly 104 times higher than that of other DNA viral genomes [28]. Thus, much like RNA and retroviruses, an intra-host virus population, referred to as the viral quasispecies, arises during HBV infections. This quasispecies consists of major, intermediate and minor variants which occur at frequencies of >20, 5-20, and <5 %, respectively [13].

As selection occurs on the entire population [3], and progresses over the course of infection [16], the population dynamics cannot be understood from the fittest strain alone; necessitating the study of viral populations at the intra-host level, in addition to the inter-host level. This has previously been limited by the inability to directly sequence the full quasispecies using conventional methods, however, the advent of next generation sequencing (NGS), which allows for the mass parallel sequencing of mixed samples, has largely circumvented this. Consequently a cornucopia of tools now exist to measure quasispecies attributes [18], such as quasitools [37] or QSutils [19], or directly reconstruct a viral quasispecies e.g., k-GEM [36], ShoRAH [62], Vispa [2], ViQuas [24], QuasiRecomb [59], and QuRe [48, 47]. Of these algorithms, k-GEM, QuasiRecomb, and QuRe are implemented in Java and take similar input, allowing cross-platform scaling of quasispecies reconstruction and a direct comparison of algorithm efficacy. Furthermore, QuRe was first tested and benchmarked on actual data from HBV genotypes A and D and accurately reconstructed the viral quasispecies for HBV with improved efficacy as compared to ShoRAH [46].

Quasispecies dynamics have been studied in HBV in a myriad of ways that evaluated quasispecies and single nucleotide polymorphisms (SNPs) in relation to key treatment and disease progression outcomes such as therapeutic response or resistance to antivirals [11, 52] and seroconversion of viral proteins [12, 33, 34], among others. Several key concerns do, however, remain to be addressed. These include whether NGS can detect more SNPs than conventional methods, if quasispecies reconstructions accurately reflect known within and between host diversity, and if quasispecies reconstruction and analyses provide comparable output regardless of the specific methods used.

The aim of the present study is to characterise quasispecies dynamics across the full genomes of HBV from patients with chronic infections. Specific objectives are to (i) compare different but similar quasispecies reconstruction algorithms that are implemented in Java with regards to their fidelity in constructing similar haplotypes, (ii) compare the resolution of detecting SNPs through NGS and quasispecies reconstruction to those typically detected using traditional methods, (iii) compare the diversity of quasispecies in relation to HIV co-infection and HBV serology markers, and (iv) provide clarity on the phylogenetic relatedness of these strains.

## Methods

### Specimen selection, serology, and ethics approval

The specimens used in the present study (n = 15) were collected as part of a larger urban hospital cohort of HBV patients from 2007-2011 by Mayaphi et al. [38, 39] and were selected for cross-sectional full genome quasispecies analyses. This follows after presenting as atypical outliers in a phylogenetic analysis of typical HBV genotypes A1 and D from South Africa using the pre-core/core and surface genes [39]. Further samples were included of HBV genotype E from a paediatric outbreak from the same hospital [45]. Sample details were deposited to the National Centre for Biotechnology Information (NCBI) BioSample database with links to the relevant BioProject (see data availability section). These specimens represented both HIV positive (n = 10) as well as HIV negative (n = 5) patients for which other serological data related to HBV infection, including antigen and antibody tests conducted by enzyme linked immunosorbent assays (ELISA), were available (Table 1). Protocol approval was obtained from the Research Protocol Committee and ethics approval from the Research Ethics Committee of the Faculty of Health Sciences, University of Pretoria (numbers: UP35/2007 and S137/2012).

**Table 1.** Summary of serology for individuals (n = 15).

| ID | Primary markers |  |  | Secondary markers |  | Molecular |  |  |
| --- | --- | --- | --- | --- | --- | --- | --- | --- |
|  | HBsAg | Anti-HBs | Anti-HBc | HBeAg | Anti-HBe | Liver ALT (U/L) | Viral load (IU/mL) | Genotype |
| 3269* | + | - | + | - | - | 21 | 9.5 x 10<sup>1</sup> | A1 |
| 3274* | + | - | + | + | - | 60 | 10.0 x 10<sup>6</sup> | A1 |
| 3319* | - | - | + | - | - | 35 | 12.7 x 10<sup>1</sup> | A1 |
| 3358* | + | - | + | - | - | NA | NA | A1 |
| 3658* | + | - | + | - | + | 29 | 17.0 x 10<sup>6</sup> | A1 |
| 3768* | - | - | + | - | - | NA | NA | A1 |
| 3791* | + | - | + | + | - | 58 | >110.0 x 10<sup>6</sup> | A1 |
| 4070* | + | - | + | + | - | 46 | >110.0 x 10<sup>6</sup> | A1 |
| 4312* | + | - | + | + | - | 61 | 7.0 x 10<sup>6</sup> | A1 |
| LA05* | + | - | + | - | + | NA | NA | A1 |
| N005 | + | - | + | - | + | 21 | 2.0 x 10<sup>3</sup> | A1 |
| N011 | + | - | + | + | - | 28 | >110.0 x 10<sup>6</sup> | D4 |
| N060 | + | - | + | - | + | 44 | 15.7 x 10<sup>1</sup> | A1 |
| N199 | - | - | + | - | - | 30 | 9.9 x 10<sup>1</sup> | A1 |
| PO04 | - | - | + | - | - | NA | NA | E |

\* HIV positive samples

### Full genome PCR amplification and NGS

Total viral genomic DNA was extracted from 100 μL stored plasma samples with the QIAamp MinElute Virus Spin Kit (Qiagen, Hilden, Germany), according to manufacturers’ instructions, and the extracted DNA eluted to a final volume of 20 μL. DNA, of the full genomes of HBV (±3221bp), was amplified by long-range polymerase chain reaction (PCR) as previously described by Günther et al. [21, 22] with minor modifications [15]. The PCR primers that were used were as follows: Forward, P1 (1821-1841), 5’-CTT TTT CAC CTC TGC CTA ATC A-3’; Reverse, P2 (1825–1806), 5’-AAA AAG TTG CAT GGT GCT GG-3’, or P2A1 (1825-1806), 5’-AAA AAG TTG CAT GAT GAT GG-3’. Success of amplification was monitored by separating reactions by electrophoresis on a 2 percent TBE-agarose gel. PCR products were purified by means of the DNA Clean and Concentrator-25 spin column kit (Zymo Research Corp., Irvine, California, USA) according to manufacturers’ instructions, prior to sequencing. Amplified and purified DNA was sent to Inqaba Biotech Inc. (Hatfield, Pretoria, South Africa) for sequencing on the Illumina MiSeq sequencing platform (Illumina (Pty) Ltd., San Diego, California, USA).

### NGS quality control, filtering, and read assembly

The FASTQ sequence files were analysed for quality (Fig. S1) and assembled to a reference in Geneious Prime 2022 (www.geneious.com). Read quality was assessed for each file to determine quality score distributions, read length distribution, overrepresented sequences and k-MERs. Paired read files were set and interleafed into a single file per specimen and were directly used for mapping and SNP calling within Geneious. Coverage was calculated for individual concatenated paired read files using the Lander-Waterman equation where coverage (C) is calculated by multiplying read length (L) by the total read number (N) and divided by the size in base-pairs of the haploid genome (G).

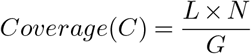

For purposes of quasispecies reconstruction the files were down-sampled to account for between specimen variation in coverage by concatenating the paired end reads using BB-Merge [9] in Geneious under standard parameter settings, allowing for the inclusion of reads with Phred-scale quality scores higher than twenty and set to remove unpaired reads. Phred-scale quality scores (Q) are computed with the following equation by taking the logarithm of the base-call error probabilities (P).

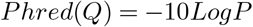

Duplicate reads were subsequently removed from the concatenated read files with Dedupe V.38.37 function implemented in Geneious with a k-MER threshold of thirty. Hereafter the remaining reads were filtered by size by extracting those with a minimum length of 100 bp. The interleafed read files were mapped to references AB048703 (genotype D) for specimen N011, DQ060823 (genotype E) for PO04 and AY233283 (genotype A) for the remainder.

### Full genome quasispecies reconstruction

The viral quasispecies was reconstructed for each specimen (n = 10) where sufficient read-depth or coverage was available by the three tested Java executable algorithms: QuRe V.0.99971 [48, 47], QuasiRecomb V.1.2 [3] and k-GEM V.0.3.1 [36].

Each algorithm was executed from the command prompt interface with the Java developers’ kit V.1.7.0-25. The merged, de-duplicated and filtered read files in FASTA format were used as input data for QuRe, along with the appropriate references used in prior mapping. In the case of QuRe, a dictionary was built from the parsed reference and read files from which a quasi-random alignment score distribution was calculated. This was followed by an internal alignment with Jaligner and the removal of reads with an alignment p-value>0.01. The reconstruction algorithm of QuRe was run in three phases: a fixed-size sliding window overlap, random overlaps, and assessment of the best a-posteriori overlap set. Thereafter the core reconstruction was run followed by the final clustering of the initial variants based on a random search and Bayesian Information Criterion (BIC) determination. For QuasiRecomb and k-GEM, the reduced reads mapped to references (in SAM format) were used as input data. In each instance, the input files were first parsed prior to model training and reconstruction based on the best model. QuasiRecomb was first run under default settings with automatic model selection and a range of one to five generators. This output was executed a second time with the refinement parameter. In k-GEM the reconstructions were run with a model selection between two and five generators. All results were standardized by down-sampling and using the same references across all experiments and algorithms.

### Full genome phylogenetic analysis

Appropriate reference sequences, retrieved from GenBank as a population model, were imported into the same file as the full quasispecies for all study specimens from the three reconstruction methods. Multiple- and pair-wise alignment of the sequence file was done using MAFFT V.7.273 [27], with automatic model detection, and the resulting aligned sequences saved in PHYLIP format. A model test was conducted in MEGA X [30] to establish which model best describes the phylogeny of the data based on the BIC (Table S1). Phylogenetic analysis was done by means of the maximum likelihood method, as implemented in PhyML V.3.1 [20]. The analysis was performed under the general time reversible (GTR + I + R) model, with gamma-distributed rate variation across sites and a proportion of invariable sites and executed a second time for bootstrap analysis with 1000 replicates. The resulting tree was viewed in Figtree V.1.6.6 [49] from which it was exported in NEWICK format. Hereafter, the maximum likelihood tree was imported into MEGA for final editing.

### Quasispecies diversity estimates and statistical analysis

Measures of quasispecies diversity [18] were measured using quasitools V.0.7.0 [37] implemented in PYTHON V.3.9, for the pooled reconstructed quasispecies, as FASTA files (n = 10), and for aligned BAM files from Geneious; for samples with incomplete coverage (n = 5). Quasispecies metrics were calculated at three levels including incidence metrics (entity level), abundance metrics (molecular level), and functional metrics (incidence level). Incidence metrics included the number of haplotypes and the total number of intra-host SNPs. Abundance metrics were computed in terms of Shannon entropy and Gini-Simpson statistics. Three functional metrics, the mutation frequency (Mfe), functional attribute diversity (FAD), and sample nucleotide diversity (π), were also computed. Statistical analyses were done using R V.4.1.3 implemented in RStudio V.2022.07.0 [57]. To ascertain if NGS increases SNP detection, a model population of HBV genomes from sequence databases was created by retrieving randomly sampled full genome sequences (n = 70) of subgenotype A1 from the National Centre for Biotechnology Information (NCBI) GenBank resource [1], to be analysed for diversity in parallel to the study specimens. The median number of SNPs called from the mapped reads in Geneious, as well as the reconstructed quasispecies, were compared to the diversity observed for the population model from GenBank with non-parametric Kruskal-Wallis tests. For the quasispecies, SNPs were computed in comparison to the reference sequence used in mapping/reconstruction, in comparison to the closest matching reference from a basic local alignment search tool (BLAST) [61] search, and internally among variants of the same sample. The number of SNPs and number of reconstructed variants were also compared to categorical variables including HIV status and HBV serology attributes, with non-parametric Kruskal-Wallis tests.

## Results

### Reconstructed quasispecies and intra-host diversity

Quasispecies were reconstructed for a total of ten samples with adequate coverage using each of the three reconstruction algorithms. A summary of the quasispecies and their relative frequencies are available in Table S2. Reconstructed quasispecies from QuasiRecomb resulted in the most individual variants (n = 55), followed by QuRe (n = 28) and k-GEM (n = 22). From k-GEM at least two variants, one major and one intermediate, were reconstructed whereas QuasiRecomb and QuRe only reconstructed the main variant for half of the specimens (online supplementary data) but more than five variants for other samples. From QuRe, the quasispecies of N005 was comprised of 8 variants, 3658 and N011 had five each and 3274 and N199 each had three. From specimen N005 one major variant, two intermediate and five extremely low frequency minor variants were reconstructed. The variants of specimen 3658 had a major variant, several low intermediate variants and two minor variants. Within the quasispecies of N011 there was one major variant and four minor variants. The two samples, 3274 and N199, had three variants each. Of the quasispecies reconstructed by QuasiRecomb six specimens comprised of more than one major variant. Two samples, N005 and N199 had the most diverse quasispecies with 14 and 23 variants each. Specimen 3274 had a quasispecies comprising four variants, 3791 comprised of six, and N011 and PO04 had two each. There was an overlap between QuasiRecomb and QuRe with regards to which specimens gave multiple variants and the relative number of SNPs between them. k-GEM generally gave two variants as output, one major variant which occurred at a frequency between 70 and 85 percent and one intermediate variant. There was, however, a lower degree of variability within the quasispecies from k-GEM than by other methods. Pooled quasispecies incidence metrics of intra-host variability (Table 2) showed “within specimen” haplotypes (H) ranging from 4 to 28 variants and a median number of SNPs of 48.00±17.52 (SEM). For the abundance metrics, a Shannon entropy (Hs) ranging from 0.69 to 3.43 was detected with a corresponding Gini-Simpson statistic of approximately 0.90±0.04. The three measures of functional diversity, mutation frequency (Mfe), functional attribute diversity (FAD), and sample nucleotide diversity (π), had mean values of 0.0096±0.0029, 9.6 × 10-3, and 0.014±0.005 respectively.

**Table 2.** Summary of NGS and quasispecies metrics (within host diversity) of study specimens

| Sample ID | Coverage (C) | Phred (Q) | Incidence (Entity) | | Abundance (Molecular) | | Gini-Simpson | Functional (Incidence) | | FAD | Diversity ($\pi$) |
| --- | --- | --- | --- | --- | --- | --- | --- | --- | --- | --- | --- |
|  |  |  | Haplotypes (H) | SNPs | Shannon entropy (HS) |  |  | Mutation frequency (Mfe) |  |  |  |
| Average: | | | 11.53 | 68.13 | 1.78 | | 0.9 | $9.6 \times 10^{-3}$ | | 0.96 | 0.014 |
| Reconstructed quasiespecies: |  |  |  |  |  |  |  |  |  |  |  |
| 3274 | 81,590 | 27 | 8 | 20 | 1.91 | | 0.84 | $9.0 \times 10^{-4}$ | | 0.066 | 0.0016 |
| 3319 | 78,190 | 26 | 4 | 34 | 1.39 | | 0.75 | $2.6 \times 10^{-3}$ | | 0.047 | 0.0039 |
| 3658 | 90,714 | 24 | 8 | 34 | 1.91 | | 0.84 | $1.5 \times 10^{-3}$ | | 0.112 | 0.0027 |
| 3791 | 80,614 | 30 | 9 | 27 | 2.04 | | 0.86 | $1.0 \times 10^{-3}$ | | 0.105 | 0.0019 |
| 4070 | 67,603 | 28 | 4 | 15 | 1.39 | | 0.75 | $1.9 \times 10^{-2}$ | | 0.458 | 0.0382 |
| 4312 | 76,646 | 28 | 4 | 10 | 1.39 | | 0.75 | $1.4 \times 10^{-2}$ | | 0.335 | 0.0279 |
| N005 | 69,552 | 26 | 24 | 184 | 3.18 | | 0.96 | $2.5 \times 10^{-3}$ | | 1.958 | 0.0035 |
| N011 | 30,455 | 25 | 9 | 54 | 2.04 | | 0.86 | $2.0 \times 10^{-3}$ | | 0.201 | 0.0036 |
| N199 | 46,738 | 22 | 31 | 248 | 3.43 | | 0.97 | $2.5 \times 10^{-3}$ | | 3.942 | 0.0042 |
| PO04 | 3,572 | 27 | 4 | 9 | 1.39 | | 0.75 | $7.0 \times 10^{-4}$ | | 0.017 | 0.0014 |
| Geneious assembly: |  |  |  |  |  |  |  |  |  |  |  |
| 3269 | 32,196 | 24 | 7 | 48 | 0.72 | | 0.85 | $7.5 \times 10^{-3}$ | | 0.06 | 0.0088 |
| 3358 | 11,499 | 28 | 13 | 83 | 1.56 | | 0.76 | $3.1 \times 10^{-2}$ | | 1.29 | 0.0404 |
| 3768 | 11,564 | 27 | 10 | 82 | 0.69 | | 0.5 | $1.8 \times 10^{-2}$ | | 0.09 | 0.0475 |
| LA05 | 6,573 | 23 | 10 | 65 | 1.09 | | 0.66 | $3.3 \times 10^{-2}$ | | 0.39 | 0.056 |
| N060 | 4,729 | 23 | 28 | 109 | 2.09 | | 0.83 | $2.6 \times 10^{-2}$ | | 5.99 | 0.0323 |

Coverage (C) and quasiespecies metrics, as determined for the reconstructed quasiespecies and for the Geneious assemblies for samples with incomplete coverage. The anticipated coverage (C) per base, based on read files, was calculated with the Lander-Waterman equation while the average Phred-score (Q) was determined in Geneious. Quasiespecies metrics we computed using quasitools and are reported at three levels: (1) incidence metrics at the entity level, (2) abundance metrics at the molecular level, and (3) functional metrics of incidence. For incidence at the entity level two metrics are reported, the number of reconstructed or detected haplotypes (H) and the total number of intra-host SNPs. Abundance metrics are given by the standard Shannon entropy (HS) and Gini-Simpson statistics, while functional metrics are given as the mutation frequency (Mfe), functional attribute diversity (FAD), and sample nucleotide diversity ( $\pi$ ).

### Inter-host diversity

From the full model population file (n = 70) of sequences retrieved from GenBank, an average of 76±25 SNPs were detected between individuals. Using a smaller population model equal in size to the study samples (n = 15), a similar average of 79±21 SNPs was detected.

Summary statistics of the quantitative variables used in statistic comparisons are provided in Table 3. The average number of SNPs between the individual quasispecies and the reference used for mapping and reconstruction ranged from 70.20±11.35 for QuasiRecomb to 77.10±11.48 for k-GEM. A statistical comparison showed no significant increase in the number of SNPs detected through mapping of read files in Geneious or through quasispecies reconstruction, as compared to the SNPs detected with conventional methods, nor a significant difference between the different reconstruction methods used (Fig. 2, χ^2^= 1.64, df = 4, p-value = 0.80, η2 = 0.04). SNPs called for the quasispecies in comparison to the closest BLAST match was generally lower (45.53±12.76) and did not significantly deviate from the median of the intrahost diversity (χ^2^= 0.50, df = 1, p-value = 0.50, η2 = 0.01).

**Table 3.** Summary statistics for variables used in statistical analysis.

| *Quantitative variables:* |  |  |  |  |  |  |
| --- | --- | --- | --- | --- | --- | --- |
| Variable (x) | Total | Min (xmin) | Max (xmax) | Mean | STD ($\sigma$) | SEM (σM) |
| SNPs (Inter-host): |  |  |  |  |  |  |
| *GenBank* | 1187 | 29 | 108 | 79.13 | 21.07 | 5.44 |
| *Geneious* | 1053 | 14 | 115 | 70.2 | 34.1 | 8.81 |
| *QuRe* | 755 | 24 | 132 | 75.5 | 35.55 | 11.24 |
| *QuasiRecomb* | 702 | 21 | 125 | 70.2 | 35.91 | 11.35 |
| *k-GEM* | 771 | 21 | 127 | 77.1 | 36.31 | 11.48 |
| SNPs (Intra-host): |  |  |  |  |  |  |
| *All samples* | 1022 | 9 | 248 | 68.13 | 67.88 | 17.52 |
| Quasispecies (Haplotypes): |  |  |  |  |  |  |
| *QuRe* | 28 | 1 | 8 | 2.8 | 2.44 | 0.77 |
| *QuasiRecomb* | 54 | 1 | 23 | 5.4 | 7.38 | 2.33 |
| *k-GEM* | 22 | 1 | 5 | 2.2 | 1.03 | 0.32 |
| *All samples* | 173 | 4 | 31 | 11.53 | 8.85 | 2.28 |
| *Qualitative variables:* |  |  |  |  |  |  |
| Variable | Total | Positive | Negative | NA |  |  |
| Serology: |  |  |  |  |  |  |
| *Human Immunodeficiency Virus (HIV)* | 60 | 28 (0.47) | 17 (0.28) | 15 (0.25) |  |  |
| *HBV Surface Antigen (HBsAg)* | 60 | 32 (0.53) | 13 (0.22) | 15 (0.25) |  |  |
| *HBV Surface Antigen Antibody (Anti-HBs)* | 60 | 0 (0.00) | 45 (0.75) | 15 (0.25) |  |  |
| *HBV Core protein Antibody (Anti-HBc)* | 60 | 45 (0.75) | 0 (0.00) | 15 (0.25) |  |  |
| *HBV e protein Antigen (HBeAg)* | 60 | 20 (0.33) | 25 (0.42) | 15 (0.25) |  |  |
| *HBV e protein Antibody (Anti-HBe)* | 60 | 10 (0.17) | 35 (0.58) | 15 (0.25) |  |  |
| HBV Genotype: |  |  |  |  |  |  |
| *Genotype A* | 52 |  |  |  |  |  |
| *Genotype D* | 4 |  |  |  |  |  |
| *Genotype E* | 4 |  |  |  |  |  |

Basic attributes of the variables (x) used in subsequent statistical analyses are summarized by ways of descriptive statistics. This included the quantitative variables, such as SNP and quasispecies numbers from each method, as well as qualitative or categorical variables such as HIV status or HBV related serological markers. For quantitative variables the sample size (n), total, minimum (xmin), maximum (xmax), Mean (x), standard deviation ( $\sigma$ ), and standard error of the mean ( $\sigma_M$ ) is reported. For qualitative variables the actual numbers as well as the frequencies are reported.

**Fig. 1.**
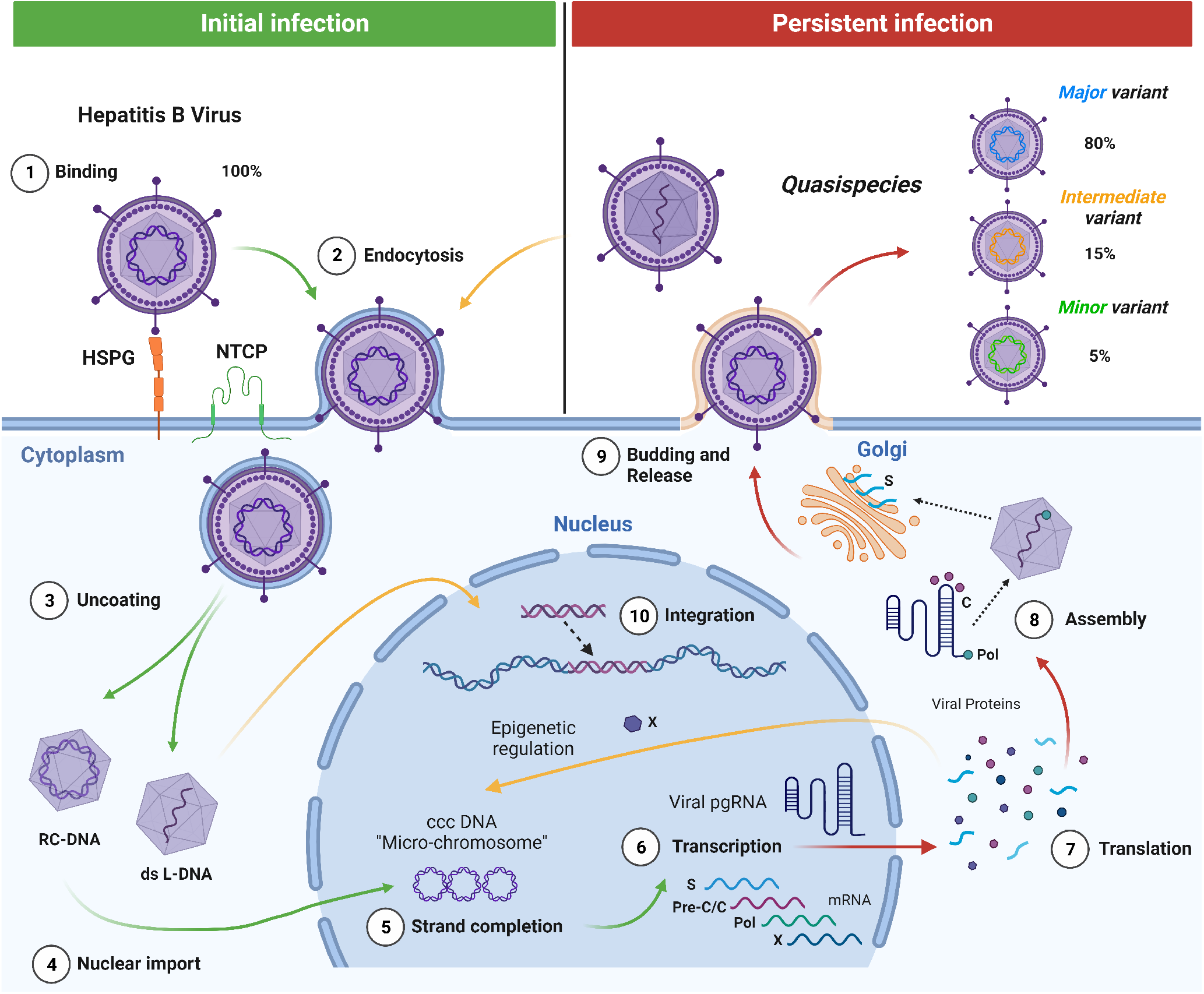
Life cycle and quasispecies formation of the hepatitis B virus. (1) Upon initial infection the virus proteins interact with membrane receptors on hepatocytes such as heparan sulphate proteoglycan (HSPG) and Na+-taurocholate co-transporting polypeptide (NTCP) to fuse with the membrane (2) and enter the host cell as a coated virion. After uncoating (3), the virus replicates by nuclear import of the relaxed circular DNA (RC-DNA) viral genome prior (4) which undergoes strand completion (5) to form covalently closed circular DNA (cccDNA) mini-chromosomes. Viral mRNA is then transcribed (6) to form the viral pre-genomic RNA (pgRNA) and several primary transcripts for the translation of viral particles and proteins (7). These proteins and particles are triggered to assemble (8) the virion through the encapsidation signal of the pgRNA. The X protein acts as an epigenetic regulator of transcription by interfacing with the micro-chromosome in the nucleus. In the assembled virion, pgRNA is reverse transcribed by viral polymerase to form the partially double-stranded DNA genome. Newly formed virions, inside specialised vesicles from the Golgi, then bud (9) through the cell membrane and are released into the blood stream. Mutations, caused by the low replication fidelity and lack of proofreading from viral polymerases, results in the accumulation of a viral quasispecies or mutant cloud in the host with several major, intermediate, and minor variants in circulation. Apical reinfection with virions that have double-stranded linear DNA (ds L-DNA) instead of RC-DNA may result in the integration (10) of viral DNA into the host genome. (image created in BioRender.com)

**Fig. 2.**
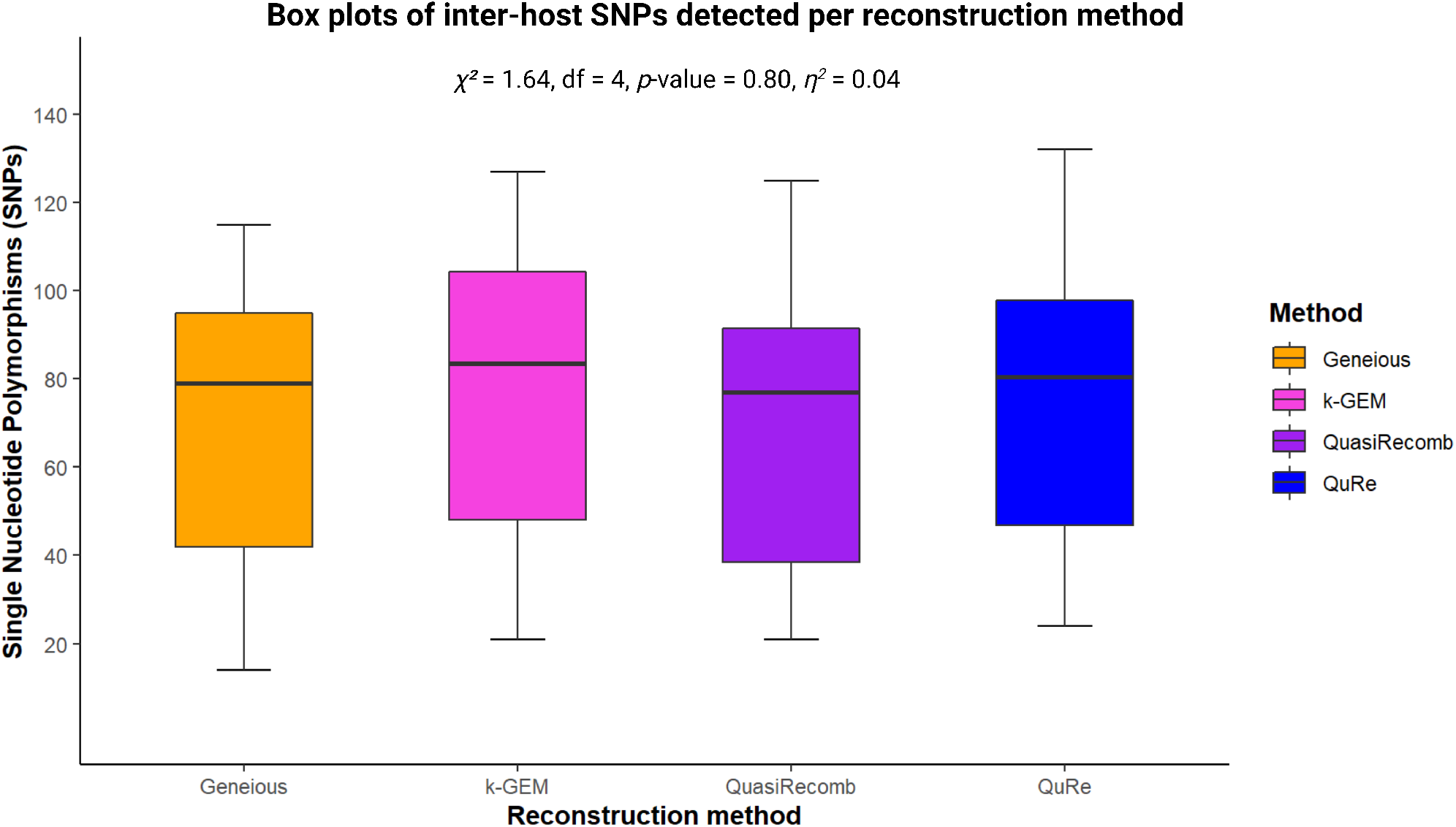
Box plots indicating the distributions of detected SNPs from assembled read files in Geneious or the quasispecies constructed through the three algorithms: QuRe, QuasiRecomb and k-GEM. The resulting SNPs were found to be similar to the typical pair-wise difference obtained for between individual variation in strains on GenBank and did not differ significantly between methods used (p-value > 0.05, 95% Confidence Interval (CI)).

### Diversity in relation to serology

Quasispecies diversity attributes including haplotype number and inter-host SNPs were compared to six serology attributes (summarised in Table 3) including HIV co-infection status and serology markers for HBV from ELISA assays. All samples were positive for HBV core protein antibody (Anti-HBc) and negative for HBV surface protein antibody (Anti-HBs). For the four remaining markers a significant relationship was detected between HIV status and haplotype number while a significant relationship was detected between SNPs and both HBV surface antigen (HBsAg) detection and HBV e-antigen antibody (Anti-HBe) detection. The number of reconstructed haplotypes were moderately increased in HIV negative samples (χ^2^= 4.61, df = 1, p-value<0.05, η2 = 0.06) whereas a large increase in SNP’s were observed for either HBV marker (Table 4, HBsAg: χ^2^= 11.36, df = 1, p-value<0.01, η2 = 0.18; Anti-HBe: χ^2^= 14.84, df = 1, p-value<0.01, η2 = 0.23). These observations were evident even at a lower sampling rate due to high effect sizes, as measured by Eta-squared (η2) and the adequately powered (β-value>0.8) non-parametric statistical analyses, tested for in post-hoc analyses.

**Table 4.** Summary of statistical tests for which a significant relationship was found.

| Comparison | df | $\chi^2$ | p-value | $\eta^2$ | Effect size |
| --- | --- | --- | --- | --- | --- |
| Quasispecies (H) vs. HIV status | 1 | 4.61 | 0.03* | 0.06 | Moderate |
| SNPs (inter) vs. HBsAg | 1 | 11.36 | <0.01** | 0.18 | Large |
| SNPs (inter) vs. Anti-HBe | 1 | 14.84 | <0.01** | 0.23 | Large |

Results from statistical analyses comparing the association between specific variables that rendered significant results. Three comparisons rendered significant results, the first between quasispecies number and the latter between the number of SNPs and two of the measured serological attributes (HBsAg and Anti-HBe). In each instance the degrees of freedom (df), Chi-squared ( $\chi^2$ ) test statistic, p-value, and Eta-squared ( $\eta^2$ ) is reported; along with the interpretation for the effect size based on the $\eta^2$ . (Significance: \*p<0.05 and \*\*p<0.01)

### Quasispecies phylogenetic relatedness

The phylogenetic relatedness of the quasispecies generated by different methods in relation to each other, as well as relevant references from the GenBank population model, is illustrated in Fig. 3. The Quasispecies generated by each method clustered together as a single clade for each individual specimen; apart from 3319.1 QuRe which branched within the clade of 3791. Specimens 4070 and 4312 partitioned with references of Asian origin within an Asian A1 branch. Closely, but separate, to that clade was the quasispecies for N199. Two samples, 3791 and 3319, partitioned with typical African A1 sequences derived from South Africa. The remaining A1 specimens partitioned in clades between the Asian and African genotype A1 branches. N005 partitioned with reference AY233290 from South Africa while specimen 3274 partitioned to form a clade with AY233287, and 3658 branched with AY233284. Specimens N011 and PO04 grouped with the genotype D and E references and were used to root the tree.

**Fig. 3.**
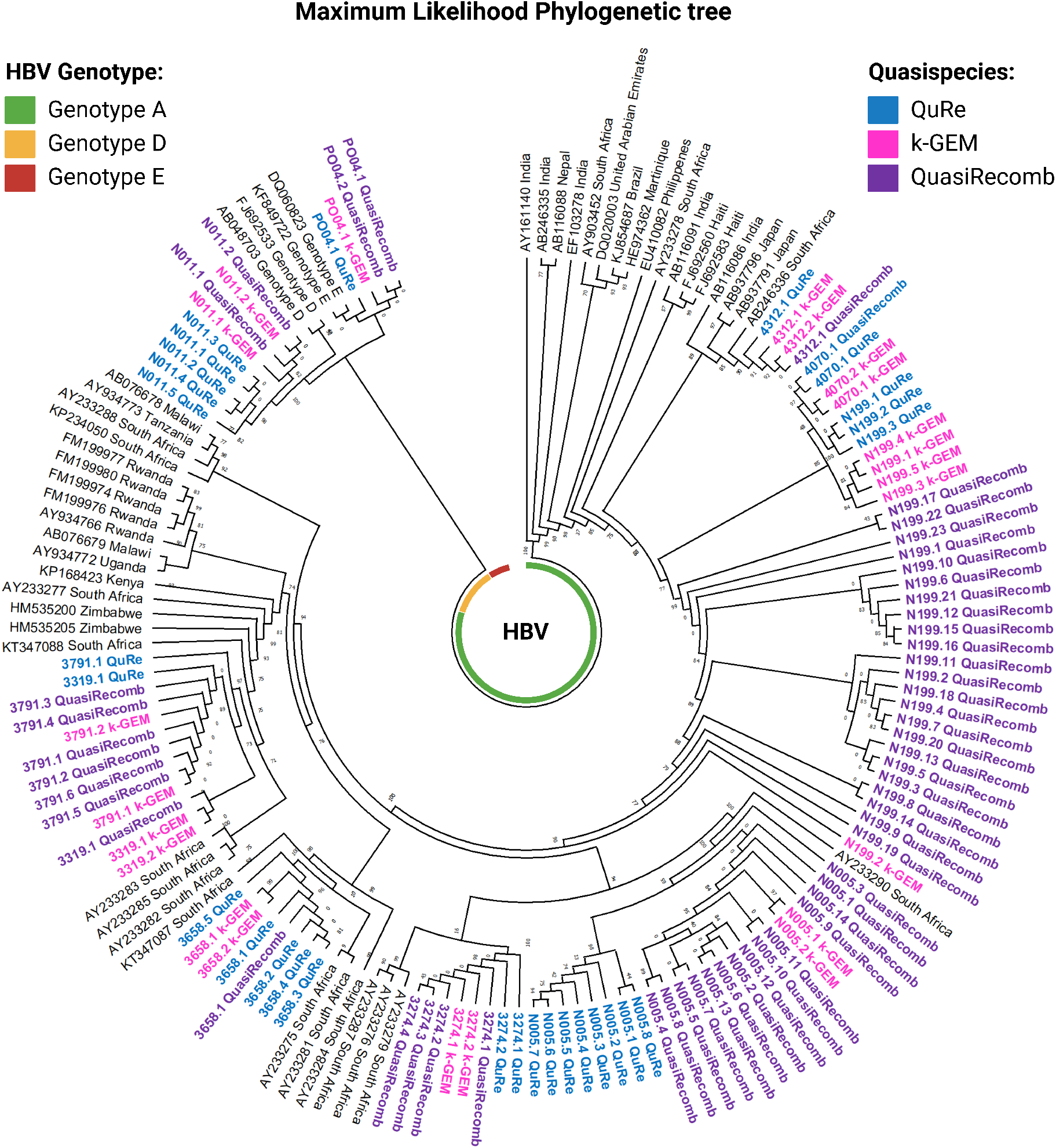
Phylogenetic tree, based on full genomes, constructed using the maximum likelihood (ML) method showing the relatedness of different reconstructed variants to established genotypes as well as between hosts differences. Quasispecies generated for the same sample by QuRe (blue), QuasiRecomb (purple), or k-GEM (pink) tended to cluster together with high bootstrap support. Most samples partitioned with genotype A references (green) while two samples, N011 and PO04, partitioned with genotypes D (yellow) and E (red) respectively.

## Discussion

Quasispecies were successfully reconstructed for several HBV specimens based on NGS sequence data with full genome coverage. Most samples reconstructed a single majority variant while many other samples had a spectrum of viral quasispecies occurring at frequencies from major variants (>20% abundance) to very low minority variants (<2% abundance). The number of detected SNPs for the quasispecies, in relation to a mapping reference, did not differ significantly from the number observed between full genome isolates of the same genotype as retrieved from online databases. Furthermore, the number of SNPs detected from each algorithm were similar to the differences between closely related isolates, such as subgenotypes, identified through BLAST searches and did not differ significantly between reconstruction methods. Phylogenetic analysis revealed that most of the isolates partition in quasispecies specific clades within HBV genotype A1. The observation that one sample partitioned within the clade of another was likely due to similarities between locally transmitted strains [31], co-infection with two strains of subgenotype A1 [4], or similar evolutionary pressures resulting in convergent evolution [58]. The outliers identified by Mayaphi et al. [39] were more closely related to Asian isolates of this genotype than characteristic South African isolates, similar to previous observations in related isolates [7]. This indicates, for the first time, that quasispecies reconstructions faithfully model known or anticipated variation between full genomes of HBV and reflect the natural evolutionary constraints of this virus.

In comparing diversity measures to HIV status, increased quasispecies diversity, as estimated by haplotype number, was observed for HIV negative specimens. Thus, the hypothesis that HIV co-infection would result in a more diverse quasispecies [43], due to a lack of immune pressure, was incorrect in this case. This would indicate that immune pressures may directly drive or enhance diversity in viral quasispecies. A positive correlation was also found for an increase in SNPs and the seroconversion, or the appearance of specific antigens and antibodies during infection, of two key HBV markers, HBsAg and Anti-HBe. This supports the observation that seroconversion is highly correlated to an increase in viral quasispecies diversity [12, 33, 34]. Conversely, because these markers change predictably over time since initial infection [29], the increase in diversity may also be related to the time period since exposure and the establishment of a chronic infection given the cross-sectional nature of this study. Other measures of quasispecies diversity were consistent with previous findings for HBV [16, 52], including a mutation frequency similar to those observed for RNA viruses but significantly higher than those typically observed for DNA viruses [28].

At present, when studying viral quasispecies, scientists are inundated with an abundance of tools for the reconstruction of quasispecies with little information available on the accuracy and applicability of a specific method to their study species. This is further confounded by the fact that many tools were initially benchmarked using simulated datasets rather than empirical data. Our comparison of the effect of different methods on the outcome revealed that, although the number of variants and their frequencies may differ between methods, the resolution in terms of SNPs detected remains constant between methods applied. Additionally, the consistency between SNPs detected from read assemblies versus reconstructed quasispecies, indicates that the actual diversity remains conserved even when more rigorous or stringent filtering methods are applied in reconstructing individual genomes. This does, however, indicate that—beyond having individual genomes for phylogenetic analyses—little additional information is to be gained from reconstructing the individual variants as many quasispecies metrics can also be estimated from alignments alone [37].

Next generation sequencing promises to be of great use in the study of viral evolution as applied to both foundational as well as clinical research. Clinical research has benefitted from the more sensitive detection of pathogens, initial drug resistance screening, and therapeutic monitoring [13], as has been done for several viruses including HBV [41], usually through studies of the polymerase gene. For foundational and applied virology, this study illustrate that NGS enables the detection and sequencing of the entire viral quasispecies of HBV. This included minor variant populations for which the full genome was assembled as opposed to only the polymerase gene, similar to studies of HIV [60], influenza A [25], human rhinovirus [56], and herpes simplex virus 1 [54]. This allows the detailed study of differential mutation rates and evolutionary processes across the genome and in different genomic regions through the real-time detection of novel variants as they occur within infected hosts as several host factors and virus dynamics shape the diversity and evolution of this virus [51]. This study also illustrates that NGS and quasispecies reconstruction not only greatly enhances the ability to model viral evolution but also has the potential to possibly predict future isolates. Future studies are needed to further elucidate differences in quasispecies dynamics between acute and chronic infections, as well as quasispecies dynamics in non-human hosts of HBV with different host immune responses.

## Conclusions

1. Different methods of quasispecies reconstruction, using similar approaches in their algorithms, reconstruct the same amount of diversity.
2. Intra-host diversity derived from full quasispecies analyses resembles and does not exceed diversity measures obtained from previous methods such as Sanger, matching known inter-host diversity.
3. Intra-host diversity between closely related quasispecies variants resembles the diversity observed within subgenotypes.
4. Diversity is increased in HIV-negative individuals possibly due to immune pressure.
5. Seroconversion of HBV biomarkers correlates with increases in diversity, possibly due to immune pressure.

## Acknowledgements

Images were created in BioRender.com. Parts of this research was presented at the Faculty Research day (symposium), Faculty of Health Science, University of Pretoria. This work is based on the research supported wholly/in part by the National Research Foundation of South Africa (Grant Number: 82831) and the Poliomyelitis Research Foundation (Grant number: 12/41 MSc).

## Data availability

Information on individual samples used in the study, as well as raw read files and a subset of reconstructed quasispecies were submitted to NCBI BioSample, Sequence Read Archive (SRA) and the Nucleotide collections with links to the BioProject PRJNA737147 (https://www.ncbi.nlm.nih.gov/bioproject/PRJNA737147). Accession numbers are listed in Table S3. The remainder of the reconstructed quasispecies and other data, have been made available on the Zenodo depository at: https://doi.org/10.5281/zenodo.7155147. The protocol for full genome amplification is available on Protocols.io.

---

**Table S1.** Maximum Likelihood fits of 24 different nucleotide substitution models.

| Model | Parameters | BIC | AICc | lnL | (+I) | (+G) | R | f(A) | f(T) | f(C) | f(G) |
| --- | --- | --- | --- | --- | --- | --- | --- | --- | --- | --- | --- |
| **GTR+G+I** | **147** | **55242.528** | **53738.094** | **-26721.941** | **0.37** | **0.54** | **1.58** | **0.225** | **0.284** | **0.271** | **0.22** |
| GTR+G | 146 | 55283.798 | 53789.597 | -26748.694 | n/a | 0.25 | 1.58 | 0.225 | 0.284 | 0.271 | 0.22 |
| TN93+G+I | 144 | 55474.621 | 54000.886 | -26856.342 | 0.37 | 0.53 | 1.56 | 0.225 | 0.284 | 0.271 | 0.22 |
| HKY+G+I | 143 | 55475.343 | 54011.841 | -26862.821 | 0.37 | 0.53 | 1.57 | 0.225 | 0.284 | 0.271 | 0.22 |
| T92+G+I | 141 | 55490.603 | 54047.566 | -26882.686 | 0.37 | 0.53 | 1.56 | 0.254 | 0.254 | 0.246 | 0.246 |
| K2+G+I | 140 | 55494.11 | 54061.307 | -26890.557 | 0.37 | 0.53 | 1.56 | 0.25 | 0.25 | 0.25 | 0.25 |
| TN93+G | 143 | 55512.738 | 54049.236 | -26881.518 | n/a | 0.25 | 1.55 | 0.225 | 0.284 | 0.271 | 0.22 |
| HKY+G | 142 | 55515.466 | 54062.197 | -26889 | n/a | 0.25 | 1.56 | 0.225 | 0.284 | 0.271 | 0.22 |
| T92+G | 140 | 55529.825 | 54097.022 | -26908.415 | n/a | 0.25 | 1.56 | 0.254 | 0.254 | 0.246 | 0.246 |
| K2+G | 139 | 55533.392 | 54110.821 | -26916.316 | n/a | 0.25 | 1.55 | 0.25 | 0.25 | 0.25 | 0.25 |
| GTR+I | 146 | 56336.702 | 54842.501 | -27275.146 | 0.6 | n/a | 1.54 | 0.225 | 0.284 | 0.271 | 0.22 |
| HKY+I | 142 | 56579.427 | 55126.158 | -27420.98 | 0.6 | n/a | 1.52 | 0.225 | 0.284 | 0.271 | 0.22 |
| TN93+I | 143 | 56582.777 | 55119.275 | -27416.537 | 0.6 | n/a | 1.51 | 0.225 | 0.284 | 0.271 | 0.22 |
| K2+I | 139 | 56592.994 | 55170.423 | -27446.117 | 0.6 | n/a | 1.51 | 0.25 | 0.25 | 0.25 | 0.25 |
| T92+I | 140 | 56594.708 | 55161.904 | -27440.856 | 0.6 | n/a | 1.51 | 0.254 | 0.254 | 0.246 | 0.246 |
| JC+G+I | 139 | 56675.141 | 55252.57 | -27487.191 | 0.36 | 0.53 | 0.5 | 0.25 | 0.25 | 0.25 | 0.25 |
| JC+G | 138 | 56711.011 | 55298.673 | -27511.243 | n/a | 0.25 | 0.5 | 0.25 | 0.25 | 0.25 | 0.25 |
| GTR | 145 | 60471.23 | 58987.262 | -29348.528 | n/a | n/a | 1.45 | 0.225 | 0.284 | 0.271 | 0.22 |
| TN93 | 142 | 60794.056 | 59340.787 | -29528.295 | n/a | n/a | 1.46 | 0.225 | 0.284 | 0.271 | 0.22 |
| HKY | 141 | 60799.445 | 59356.409 | -29537.107 | n/a | n/a | 1.46 | 0.225 | 0.284 | 0.271 | 0.22 |
| K2 | 138 | 60805.288 | 59392.95 | -29558.382 | n/a | n/a | 1.46 | 0.25 | 0.25 | 0.25 | 0.25 |
| T92 | 139 | 60812.433 | 59389.863 | -29555.837 | n/a | n/a | 1.46 | 0.254 | 0.254 | 0.246 | 0.246 |
| JC | 137 | 61923.887 | 60521.782 | -30123.799 | n/a | n/a | 0.5 | 0.25 | 0.25 | 0.25 | 0.25 |
| JC+I | 138 | 61936.031 | 60523.694 | -30123.754 | 0 | n/a | 0.5 | 0.25 | 0.25 | 0.25 | 0.25 |

Models with the lowest BIC scores (Bayesian Information Criterion) are considered to describe the substitution pattern the best. For each model, AICc value (Akaike Information Criterion, corrected), Maximum Likelihood value (lnL), and the number of parameters (including branch lengths) are also presented [40]. Non-uniformity of evolutionary rates among sites may be modeled by using a discrete Gamma distribution (+G) with 5 rate categories and by assuming that a certain fraction of sites are evolutionary invariable (+I). Whenever applicable, estimates of gamma shape parameter and/or the estimated fraction of invariant sites are shown. Assumed or estimated values of transition/transversion bias (R) are shown for each model, as well. They are followed by nucleotide frequencies (f) for each nucleotide pair. For estimating ML values, a tree topology was automatically computed. The analysis involved 70 nucleotide sequences. Codon positions included were 1st+2nd+3rd+Noncoding. All positions containing gaps and missing data were eliminated. There were a total of 2943 positions in the final dataset. Evolutionary analyses were conducted in MEGA [30].

**Table S2.**
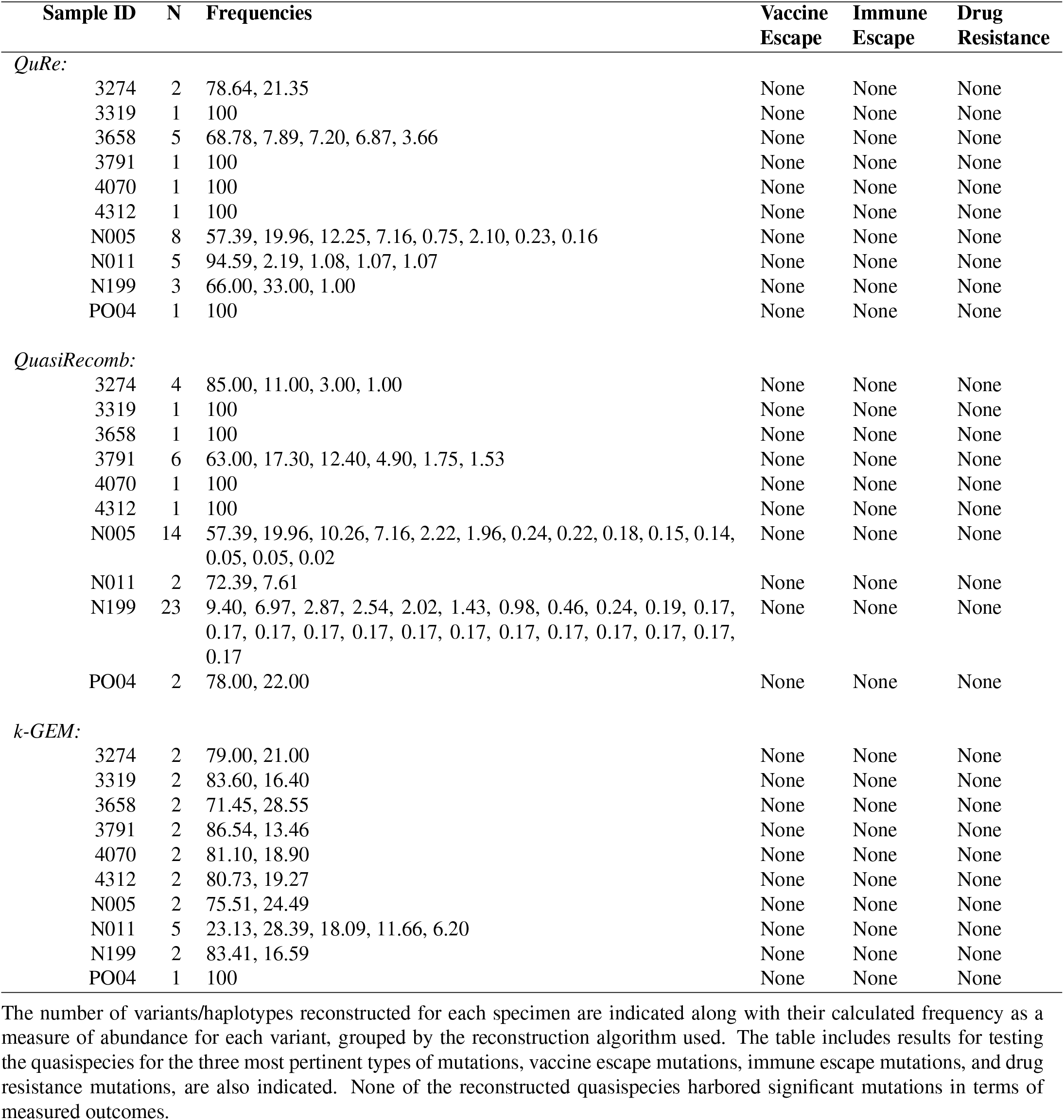
Summary of the number of variants/haplotypes reconstructed from each method along with their computed frequency.

**Table S3.** Summary of NCBI accession numbers for each study sample and associated data.

| Study ID | BioSample | Sequence Read Archive (SRA) | GenBank (Nucleotide)* | BioProject |
| --- | --- | --- | --- | --- |
| 3269 | SAMN19694634 | SRX11143873 | NA | PRJNA737147 |
| 3274 | SAMN19694639 | SRX11143872 | KF922429-KF922430 | PRJNA737147 |
| 3319 | SAMN19694636 | SRX11143879 | KF922422-KF922423 | PRJNA737147 |
| 3358 | SAMN19696888 | SRX11143880 | NA | PRJNA737147 |
| 3658 | SAMN19694641 | SRX11143881 | KF922433-KF922434 | PRJNA737147 |
| 3768 | SAMN19696889 | SRX11143882 | NA | PRJNA737147 |
| 3791 | SAMN19694632 | SRX11143883 | KF922406-KF922409 | PRJNA737147 |
| 4070 | SAMN19694637 | SRX11143884 | KF922424-KF922425 | PRJNA737147 |
| 4312 | SAMN19694638 | SRX11143885 | KF922426-KF922428 | PRJNA737147 |
| LA05 | SAMN19694648 | SRX11143886 | NA | PRJNA737147 |
| N005 | SAMN19694635 | SRX11143874 | KF922414-KF922421 | PRJNA737147 |
| N011 | SAMN19694640 | SRX11143875 | KF922432 | PRJNA737147 |
| N060 | SAMN19694646 | SRX11143876 | NA | PRJNA737147 |
| N199 | SAMN19694633 | SRX11143877 | KF922410-KF922413 | PRJNA737147 |
| PO04 | SAMN19696890 | SRX11143878 | KF922438-KF922439 | PRJNA737147 |

\* Based on original quasispecies reconstructed with QuRe in Le Clercq [14]
Information on the origin and biographic details for individual samples were submitted to the BioSample database. The associated MiSeq sequencing data was submitted to the SRA database and linked to the relevant samples. Preliminary quasispecies reconstructed with QuRe, reported in Le Clercq [14], were submitted to the core collection of Genbank Nucleotide database. All project data is linked to the registered BioProject.

**Fig. S1.**
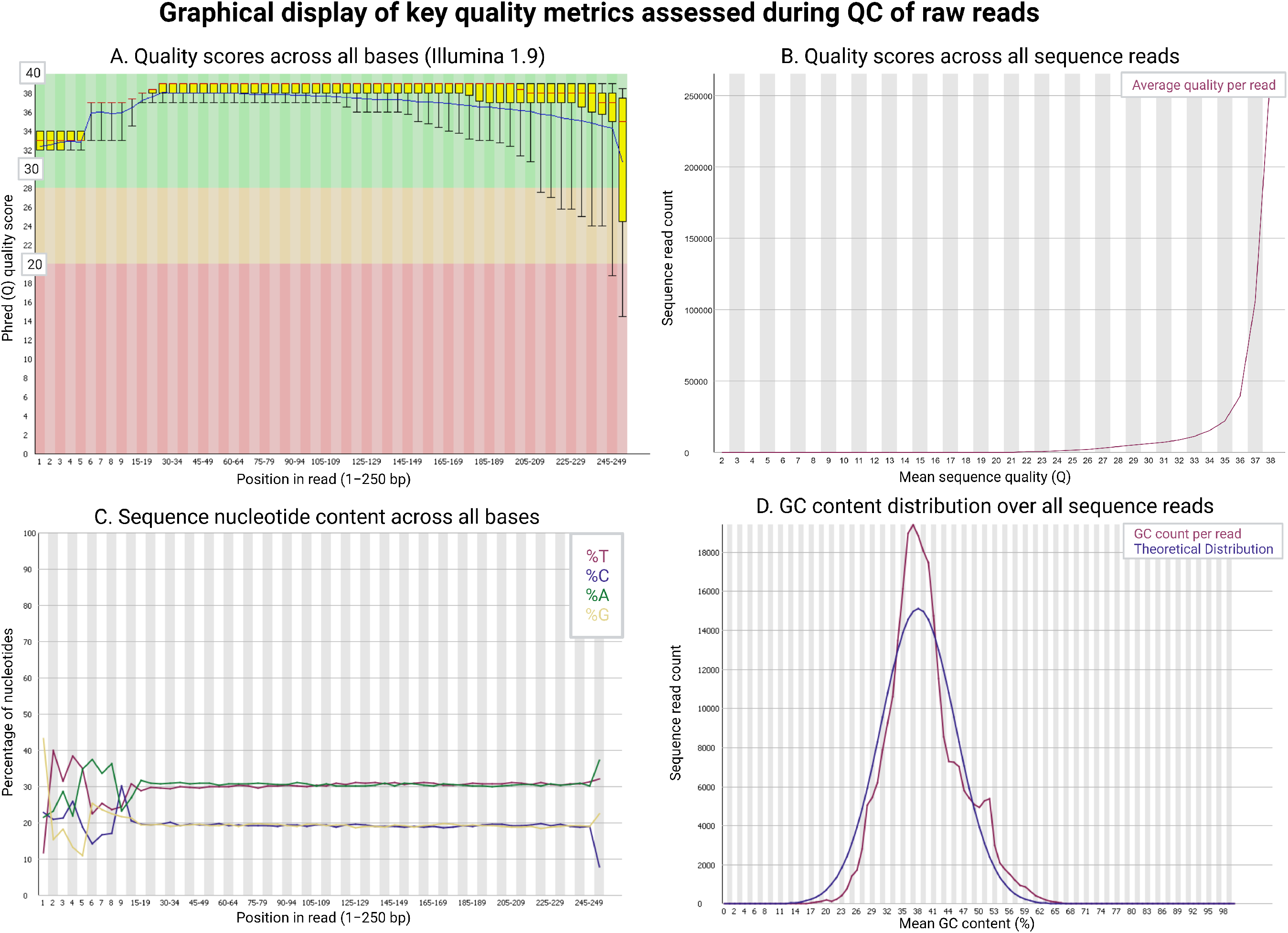
Graphical display of key metrics assessed during the quality control (QC) of read files. A) Plot of average quality scores per base from position 1-250 bp for individual reads. B) Plot of quality scores for reads based on average quality score per read. C) Plot of positional nucleotide composition (%) for reads from position 1-250 bp. D) Comparison of actual versus theoretical distribution of GC content (%).

